# Predicting Latent Links from Incomplete Network Data Using Exponential Random Graph Model with Outcome Misclassification

**DOI:** 10.1101/852798

**Authors:** Qiong Wu, Zhen Zhang, James Waltz, Tianzhou Ma, Donald Milton, Shuo Chen

**Affiliations:** Department of Mathematics, University of Maryland, College Park, MD; School of Business, Towson University, Towson, MD; Maryland Psychiatric Research Center, Department of Psychiatry, University of Maryland, Baltimore, MD; Department of Epidemiology and Biostatistics, University of Maryland, College Park, MD; Maryland Institute for Applied Environmental Health, University of Maryland, College Park, MD; Division of Biostatistics and Bioinformatics, School of Medicine, University of Maryland, Baltimore, MD

**Keywords:** link prediction, ERGM, outcome misclassification, brain network, social network

## Abstract

Link prediction is a fundamental problem in network analysis. In a complex network, links can be unreported and/or under detection limits due to heterogeneous noises and technical challenges during data collection. The incomplete network data can lead to an inaccurate inference of network based data analysis. We propose a new link prediction model that builds on the exponential random graph model (ERGM) by considering latent links as misclassified binary outcomes. We develop new algorithms to optimize model parameters and yield robust predictions of unobserved links. The new method is applied to a partially observed social network data and incomplete brain network data. The results demonstrate that our method outperforms the existing latent-contact prediction methods.

## 1. Introduction

Network data has become increasingly important to study complex problems, for example, to understand the role of contact patterns in epidemics through social network analysis (Stattner and Vidot, 2011) and interactions between neuron populations in the human brain (Simpson et al., 2011, Simpson et al., 2019). A network can be represented by a graph, where a node denotes a study unit and an edge or link indicates interactions between a pair of nodes (Zhao et al., 2017). In practice, network data are often incomplete because some links are unreported/undetectable due to various noises and technical limitations during data collection. For example, the self-reported data collection procedure for social networks is subject to the “recall bias” and the chance of a subject recollecting all his/her contacts is very small. In addition, the number of contacts that a subject can report may be restricted to a fixed number that is less than his/her total contacts. Therefore, the social network estimated from the survey data is partially observed where many links are latent (false negatives). Similarly, brain connectome network data may miss true links due to noise and weak signal levels (Simpson and Laurienti, 2015). It has been well documented that inferences based on incomplete network data can be inaccurate (Zhao et al., 2017). To mitigate this challenge, we focus on developing link prediction models for incomplete network data.

The statistical analysis of network data (mainly social network analysis) has been an active area for decades (Goldenberg et al., 2010). Markov random graphs by Frank and Strauss (1986) firstly relaxed the dyadic independence of *p*_1_ model in Holland and Leinhardt (1981). Strauss and Ikeda (1990)’s discussion made the Markov model and its general form *p∗*model to be computationally available by an approximation based on the logistic regression model via a pseudo-likelihood method (Wasserman and Pattison, 1996). The likelihood-based social network model with the assumption of conditional independence has been well accepted because it is flexible to adjust the multivariate features of each node and connections between nodes. Exponential random graph models - ERGM (Snijders and Van Duijn, 2002) and latent space approach (Hoff et al., 2002) further improve the statistical properties of the parametric social network model. Efficient computational algorithms have been also been developed using advanced algorithms (Miller et al., 2009, Al Hasan et al., 2006, O’Madadhain et al., 2005). The parametric network models have been widely used for social network analysis. Recently, network models have also been successfully applied to complex brain connectome data analysis (Simpson et al., 2019). However, these parametric network models are built on completely observed network data (i.e. without latent links) and thus may not be directly applied to the incomplete network data for latent link prediction (Martínez et al., 2017).

Predicting latent links in incomplete networks can be considered as a binary outcome prediction problem. However, the commonly used binary outcome predictive models and machine learning methods (e.g. gradient boosting and random forest) are limited for this purpose because the outcome labels in the training data are misclassified. To address this issue, non-parametric statistical methods have been used for link prediction which takes into account the network topology and structure without the requirement of correct labeling of the training set, thus are well-suited for the partially observed network data (Zhao et al., 2017; Zhang and Chen, 2018). The non-parametric models rely on similarity-based methods assuming that the nodes are more possible to have edges with other similar nodes. In addition, local, global and quasi-local approaches are further used to describe the topology information included in computing the similarity of nodes.

In this article, we consider the unreported/undetected links as missclassified binary outcomes in the framework of statistical network analysis. The binary outcome misclassification problem has been well-studied in the statistical literature, for example, using logistic regression for epidemiological research (Lyles and Lin, 2010). The likelihood-based method under non-differential and differential outcome misclassifications assumptions can perform well to correct the estimation bias when validation data sets are provided (Magder and Hughes, 1997, Neuhaus, 1999, Carroll et al., 2006). However, the major challenge to model incomplete network data with misclassified outcome models is caused by the lack of validation data, and then neither sensitivity and specificity can be estimated without using the validation data set. In that, the likelihood function can not be correctly specified for parameter estimation (Neuhaus, 1999). In addition, the misclassification mechanism of incomplete network data is different from existing binary misclassification models in logistic regression. The commonly used assumption for a partially observed network is that the reported links are true links and not false positive, while some unreported links may be latent (Clauset et al. 2008, Rhodes and Jones, 2009, Zhao et al., 2017). Besides, the goal in the current research is to build a predictive model with high accuracy of latent link prediction, which is different from the goal of estimating effects regression coefficients in the parametric model. To address these needs, we propose a new statistical network model that integrates misclassified outcome variables into ERGM and provide efficient algorithms for accurate link prediction.

The paper is organized as follows: in section 2, we describe the proposed method and optimization algorithms, and provide theoretical results for the conclusion that the performance of our link prediction is robust and accurate given the validation data set is not available. The numerical results of both simulations and two data examples in sections 3 and 4 demonstrate that the proposed approach outperforms existing methods in various settings.

## 2. Methods

To begin, we denote a true link/contact between a pair of nodes in a network *i, j* ∈ {1, …, *n*}, *i* ≠ *j* by a binary variable *y_ij_* and let *y_ij_* = 1 indicate a positive connection and *y_ij_* = 0 otherwise. The corresponding observed link in incomplete network data is *w_ij_*. In the context of latent links (e.g. epidemiological contacts in a social network), it seems sound to assume that all observed connected links are truly connected whereas some unobserved connections are latent true links (Zhao et al., 2017). Specifically, we let a proportion of true links are misclassified as latent links that Pr(*y_ij_* = 1*|w_ij_* = 0) *>* 0 whereas all reported links are considered true contacts Pr(*y_ij_* = 1*|w_ij_* = 1) = 1. Note that this outcome misclassification mechanism is different from conventional outcome misclassification models which is reflected by our model specification in section 2.2. Our goal is to predict/recover the latent contacts from observed incomplete network data: 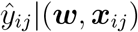 where ***w*** is the partially observed contacts and other auxiliary information such as spatial distance and co-working relationship between subjects are available in ***x****_ij_*.

### 2.1 Background

#### 2.1.1 ERGM

Let ***Y*** be a random matrix for the true contact network, ***y*** is a realization of the adjacency matrix and *Y* is the support of ***Y***. The ERGM has the form (Hunter et al., 2008):

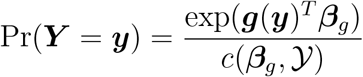

where 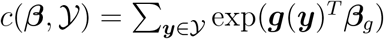 is a normalizing term with summation over all possible networks, and ***g***(***y***) is a vector of network statistics based on the adjacency matrix ***y***.

If 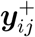 is an adjacency matrix with contact *y_ij_* = 1 and all other contacts being the same as ***y****_ij_*, and 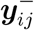 with *y_ij_* = 0. We denote the change statistics 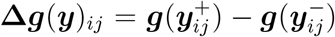 being the change of statistics from a network with *y_ij_* = 0 to *y_ij_* = 1. Then, by conditioning on the rest of network,

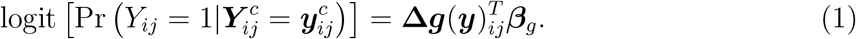

The right-hand side of (1) can be extended to 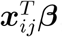, where ***x****_ij_* is a *p ×* 1 vector including the change statistics **Δ*g***(***y***)*_ij_*, node pair characteristics, and the auxiliary information of interactions, and ***β*** is the corresponding *p ×* 1 vector of parameters. In dyadic independent models, the pseudo-likelihood is equivalent to the true likelihood and provides exact estimates of parameters. In dyadic dependent ERGMs, the pseudo-likelihood may not provide valid estimates of parameters, yet, it’s still valid for link predictions because only regulative rank of links is necessary for link prediction and there is no need for calculating the normalizing terms (Shojaie, 2013). The pseudo-likelihood model is commonly used to provide a reliable and fast initial estimate of parameters in ERGM as a starting point of the MCMC algorithm (it requires to start with parameters ‘close’ to the target) (Wasserman and Pattison, 1996, Snijders and Van Duijn, 2002, Robins et al., 2007, Miller et al., 2009). Therefore, the pseudo-likelihood/likelihood function is the centerpiece of network models and to address the misclassified links.

*y_ij_* is not directly available for our incomplete network data because many links are unreported and thus latent. Clearly, simply substituting *y_ij_* with *w_ij_* in the pseudo-likelihood is invalid because *w_ij_* is subject to the misclassification errors and the equation of conditional probability on the rest of networks does not hold anymore. Therefore, we propose a new approach to incorporate outcome misclassification into ERGM.

#### 2.1.2 Misclassification model for incomplete networks

We assume the self-reported contacts *w_ij_* in the questionnaire data are observed with the non-differential single directional misclassification errors such that

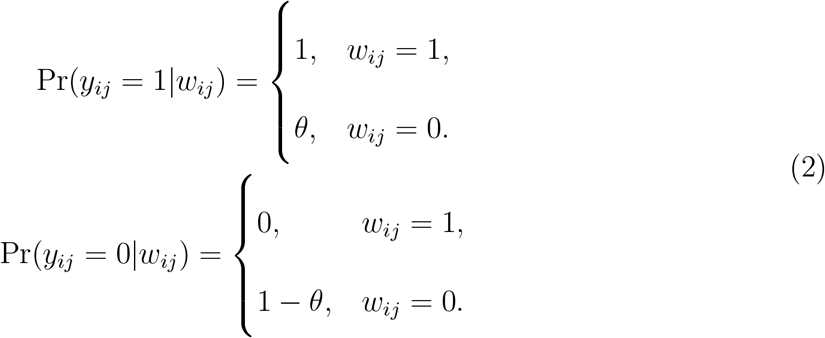

Besides, the observed nodes set are considered to be the true nodes set, such that the nodes with no corresponding edges are singletons in observed network.

We note that our outcome misclassification specification in (2) is different from the existing methods for outcome misclassification in logistic regression using the likelihood function 𝓁(***θ***; *Y, X*) = 𝓁 (***θ***; *W, X,* sensitivity, specificity) where sensitivity and specificity are estimated from the validation data (Carroll et al., 2006, Lyles et al., 2011). However, the incomplete network data often lacks validation dataset which poses difficulty to properly specify sensitivity and specificity. Thus, the likelihood function can not be optimized using the existing misclassification models. As shown by follows, our misclassification model in 2 is well suited for the misclassification mechanism of incomplete network data, and more importantly, can efficiently optimize the likelihood function for parameter estimation.

### 2.2 Expected likelihood for incomplete networks

In binary outcome misclassification models, the likelihood function can be specified based on the observed (misclassified) outcome variables (e.g. via logistic regression in Lyles and Lin, 2010). However, the distribution of misclassified outcome can be invalid for network models (e.g. ERGM). Thus, we calculate the expected value of the likelihood of ***y*** in ERGM (Reilly and Pepe, 1995): *E_θ_* (𝓁 (***β****, **y**, X*)*|****w***) based on the misclassification model (2) since 𝓁 (***β****, **w**, X*) can not be directly specified. Note that the normalizing term is omitted since we focus on link prediction (Shojaie, 2013). From (1), *y_ij_* only depends on the rest of network 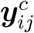 via the change statistics. We assume that the change statistics for the true network can be approximated by the observed network. The expected log likelihood function for ***y*** conditional on observed network ***w*** is:

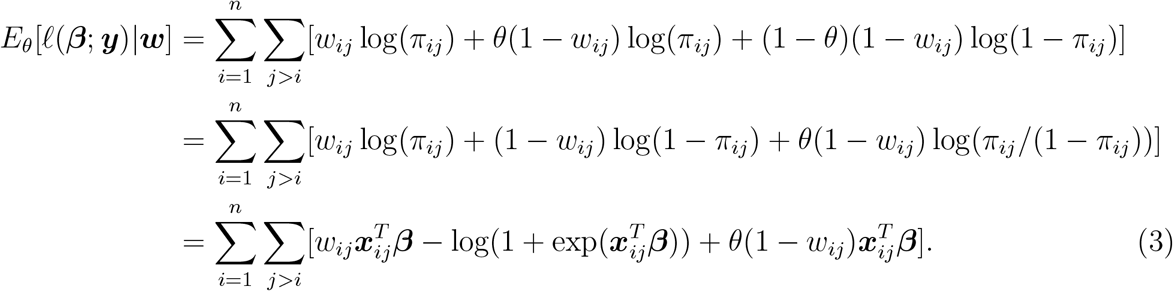

It appears that the expectation-maximization (EM) algorithm can be applied to estimate *θ* and ***β*** from (3). However, when the validation data is not available, the misclassification parameters are hard to estimate (Copas, 1988). In other words, these parameters are weakly identifiable and the EM may not converge or provide accurate estimation (Neuhaus, 1999 and Lyles et al., 2011). To address this issue, we propose a two-step procedure that estimate *θ* and ***β*** iteratively to maximize latent contact prediction accuracy.

*Step 1*: We first perform maximum likelihood estimation (MLE) of the expected log likelihood based on a given value of *θ*, which is 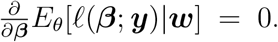. Under regularity conditions,

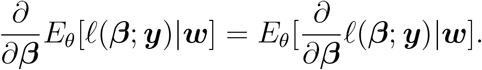

The first derivative of the expected log-likelihood is similar to the mean score function for a log-likelihood. It has been known that the MLE based on observed data can be solved by the mean score function (Reilly and Pepe, 1995).

To implement the Newton-Raphson algorithm for the estimation of ***β*** given a known value of *θ*, the first and second derivatives of the expected log-likelihood are calculated as follows:

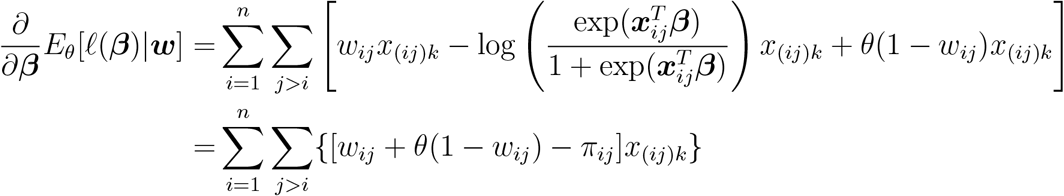

and

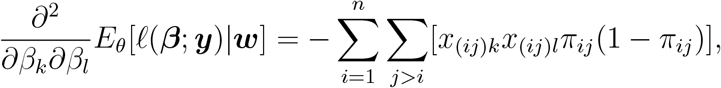

where *x*_(*ij*)*k*_ is the *k*th component of covariate vector ***x_ij_***.

Let

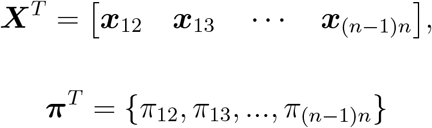

The associated matrix form for first and second order derivatives are

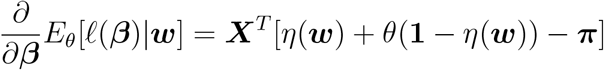

and

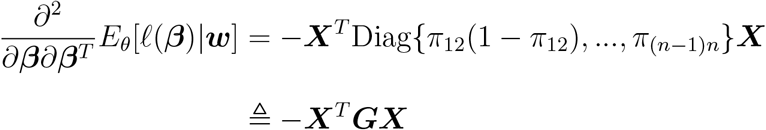

where *η*(*·*) is the half vectorization. We next update *β*^old^ by

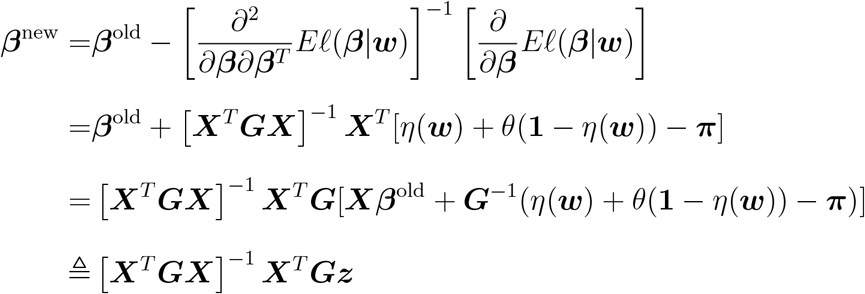

where *z* = ***Xβ***^old^ + ***G****^−^*^1^[*η*(***w***) + *θ*(**1** *−η*(***w***)) *−* ***π***]. The estimated parameters will be updated iteratively until convergence is reached. We denote the above estimated parameters as 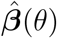 and we can accordingly calculate 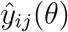 based on these parameters.

When the validation data is available and *θ* can be pre-specified by transforming sensitivity and specificity, the parameters estimated by 3 are unbiased and consistent (Carroll et al., 2006). We provide numerical validation of the unbiased and consistent estimation in section 3.3. The estimated parameters with pre-specified *θ* can be further refined by including the normalizing factor via the MCMC algorithm of ERGM (Hunter et al., 2008).

*Step 2*: With the absence of validation data, we argue that link prediction is robust although the estimate of parameter *θ* may be biased. It has been known that the estimate of *θ* is challenging without the validation data due to the issue of identifiability (Neuhaus, 1999). In step 2, we resort to an alternative pathway to optimize *θ* as a tuning parameter using the following objective function,

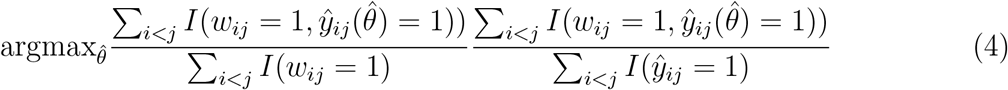

where 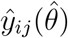 is thresholded 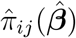 with 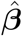 being estimated from 3 given 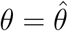. This objective function is close to the *F*_1_ score which is a balanced metric between precision and recall and is commonly used to evaluate the performance of predictive models for uneven class distributions (e.g., a large proportion of links are truly absent in a social network). The Newton-Ralphson algorithm in step one yields 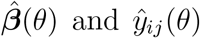 with a given *θ*. Since the range of *θ* is constrained between 0 and 1, we perform grid search to optimize 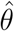 with a small incremental step (e.g. 0.001) from 0 to 1. During the grid search, we compute the value of optimization function in (4) using each value of *θ*. The optimal 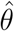 selected when maximum of the objective function (4) is achieved. Similarly, the optimal threshold *r*_0_ that 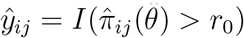 can be determined by (4).

Optimizing (4) generally yields an good estimate 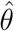 that is located within the neighborhood of true *θ*_0_ where *θ*_0_ ranges from 0 to 1. However, 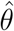 may be biased without the use of validation data which can lead to biased estimates 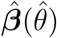(Neuhaus, 1999). Here, we argue that the biased estimates 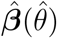 have little impact on the performance of link prediction. For example, the area of the curve (AUC) of the ROC is a commonly used metric to evaluate the performance of latent link prediction, which is approximately equal to the *c* statistic using logistic regression (Austin and Steyerberg, 2014, Harrell Jr, 2015). The *c* statistic is determined by 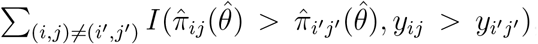, where *I* is a indicator function.

We provide both theoretical proof as follows and numerical validations in the Appendix and sections 3 and 4 to support this claim.

#### Lemma 1

For 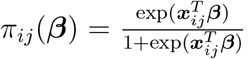, assume the estimated slope parameter is consistent up to a scalar, i.e. 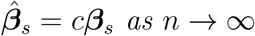 where *β* = (*β*_0_, ***β****_s_*). Then, we have as n → ∞

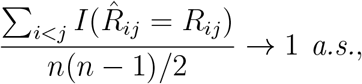

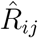 is the rank of 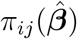, and *R_ij_* for π_ij_ (***β***).

*Proof.* From the definition,

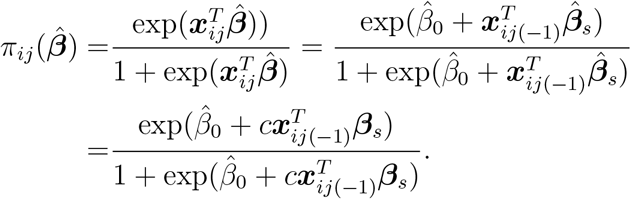

where 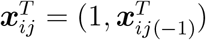

Since the logistic link function in our method is monotonically increasing, we have

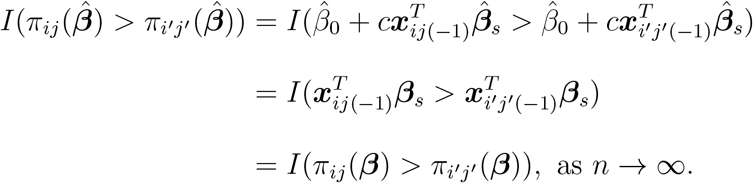

Hence,

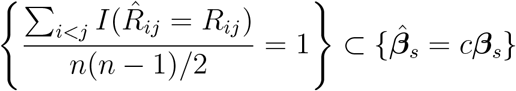

and the claim is true.

#### Theorem 1

In a network, assume true contacts given the rest of network 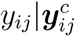 are generated independently with Bernoulli(π_ij_ (**β**)), and 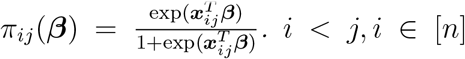. **x**_ij_ follows some distribution with (**x**_ij_, y_ij_) being exchangeable, and the conditional expection 𝔼(**bx_ij_** |**βx_ij_**) exists and is linear in **βx_ij_** for all **b** ∈ R^d^. The observed network with contacts w_ij_ are generated from model 2. θ_0_ and 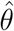 represent the true and our estimated missing rate and in general θ_0_ ≠ 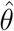. Then, as n → ∞,

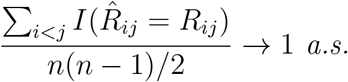

with 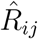 and R_ij_ being ranks of 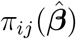 and π_ij_ (**β**), respectively. 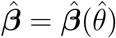 are the maximizer of 3 given missing rate 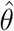.

Proof. Since the contacts in observed network are exchangeable, from the strong law of large numbers for exchangeable random variables (Kingman et al., 1978),

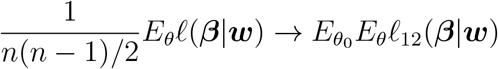

and both terms are convex in θ.

Therefore, following the results by Neuhaus (1999) and Li et al. (1989), the mis-specifcied outcome missclassification parameters (*θ*) can be considered as mis-specified link function, and 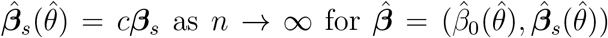 for 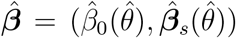 being the maximizer of the left term.

Hence, from lemma 1, the results are proved.

Therefore, the deviation of 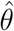 from *θ*_0_ has little impact on the proportion of the concordant pairs (i.e. 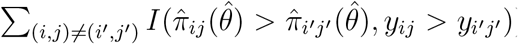), and thus does not affect the *c* statistic of the logistic regression and the AUC of ROC. The performance of our link prediction is not impacted by the variability of *θ* estimation due to the lack of validation data set, though the estimates of regression coefficients may be biased. This conclusion may partially explain that the likelihood-based social network model (with outcome misclassification adjustment) can outperform the popular models of nonparametric and machine learning techniques.

In summary, the iterative two-step procedure above provides a viable solution of link/outcome prediction for the binary regression model with the misclassified outcome but no validation data set. The proposed computational strategy is efficient and scalable to very large social networks. The complexity of the proposed method is bounded by *O*((*p* + 1)*M*), where *p* is the number of covariates and *M* is determined by the resolution of search grids for *θ* and dichotomization thresholds.

## 3. Simulation Studies

### 3.1 Synthetic data

We simulate the social network data sets by using the following models:

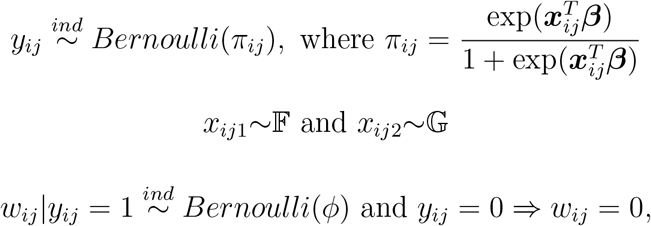

where *i, j ∈ n* denote a pair of subjects in the cohort of *n* subjects. A link *y_ij_* follows a

Bernoulli distribution with the parameter of *π_ij_*. *π_ij_* is determined by the characteristics of nodes (*x_ij_*_1_) and network topological parameters (*x_ij_*_2_). A proportion of links are latent and not reported due to recall bias and thus the partially observed network is composed of *{w_ij_}*. The single-directional misclassification mechanism is described by (2) which can be straightforwardly transformed to the sensitivity *φ* and specificity.

We simulate the networks using *n* = 150 nodes and three topological structures: 𝒯_1_ ‘random graph’, 𝒯_2_ ‘community/stochastic block’ structure with a single major community, and 𝒯_3_ ‘rich club’ (Zhou and Mondraǵon, 2004). Specifically, we let *x_ij_*_1_ *∼* Normal(*−*1, 1), *x_ij_*_2_ *∼* Exponential(2), and ***β****^t^* = (2, 1) for the random graph structure 𝒯_1_. For the community structure 𝒯_2_, we assume *x_ij_*_1_ = 1 for within community links and 0 otherwise with ***β****^t^* = (2.53*, −*2.94). In the ‘rich club’ social network data, *x_ij_*_1_ is also used to denote the topological structure and *x_ij_*_1_ = 1 for links between all ‘rich’ nodes and between ‘rich’ nodes and their ‘periphery’ nodes and with ***β****^t^* = (2.94*, −*2.94).

We repeat each setting for 100 times with different levels of recall bias (by tuning *φ* levels). For the random graph structure 𝒯_1_, we used the covariates *x_ij_*_1_*, x_ij_*_2_ in model fitting, while for community structure 𝒯_2_ and ‘rich club’ 𝒯_3_, only summarized statistics for observed network are considered, including vertex degree, number of common neighbors between vertices, shorest path between vertices, the transitivity, and so on. Via implementing model selection techniques for structure 𝒯_2_ and 𝒯_3_, the variables for vertex degree and number of common neighbors between pairs of nodes are incorporated. The latent links (*w_ij_* = 0*|y_ij_* = 1) are considered as the hold-out data and used to evaluate the performance of the link prediction by our method: statistical network model with outcome misclassification (SNOM) and the existing methods including: Neighborhood Smoothing (NS) Method by Zhang et al., 2017, Stochastic Block Model (SBM) by Airoldi et al., 2013 and Full Sum (FS) Method by Zhao et al., 2017. Note that SBM is developed for the completely observed social network data instead of partially observed data. We include this method as a reference to emphasize the importance of models addressing the latent links (misclassified outcomes) for the purpose outcome prediction. Specifically, we denote predicted edges as *y*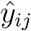, and the True Positive Rate (TPR) and False Positive Rate (FPR) were defined as:

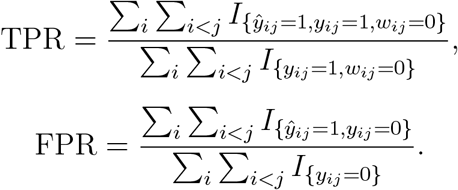

### 3.2 Results

We summarize the performance using the averaged metric of the AUC of ROC in Table 1 and Figure 1 across the 100 repetitions. We note the largest difference comes from the network data with a random graph structure, where most information is included by the characteristics of nodes and links instead of similarity of nodes and links. Therefore, our method outperforms the other machine learning and nonparametric methods because our parametric model can better leverage auxiliary covariates. For social networks with highly structured topological patterns (i.e. community and rich-club), all methods have similar AUCs when a small proportion of links are latent. SNOW seems to be more robust to a higher proportion of misreported links. The prediction SBM method is less accurate because it is developed for a fully observed network and does not account for the misclassified outcomes. Therefore, the social network models without outcome misclassification adjustment may not be applicable for partially observed social network data (Zhang et al., 2017). In summary, the simulation results demonstrate that the proposed likelihood-based social network model with outcome misclassification adjustment can provide robust and accurate predictions of latent links for various social network data.

**Figure 1:**
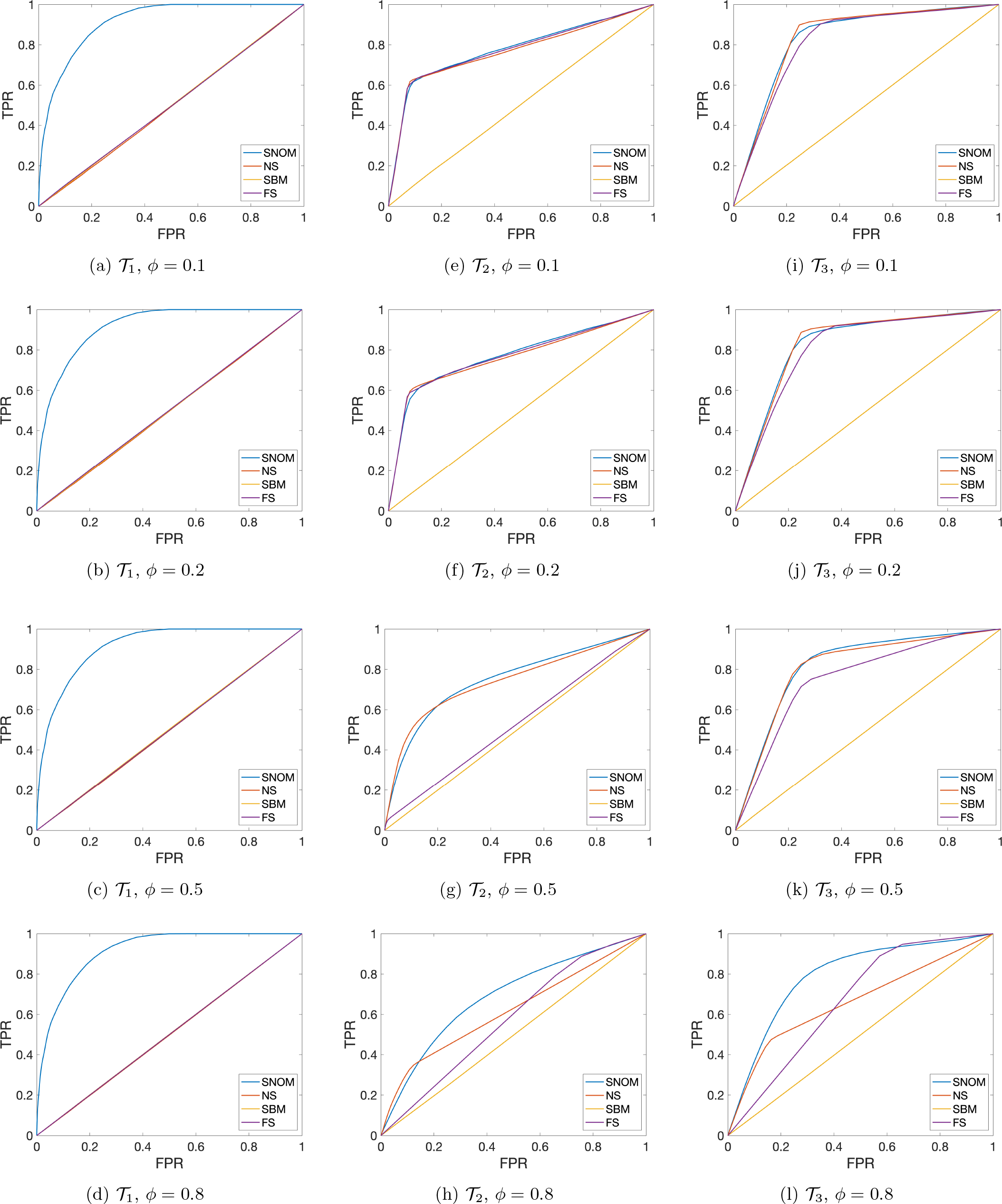
Averaged ROC curves of social networks with the structures of Random Graph (𝒯_1_), Community (𝒯_2_) and Rich Club (𝒯_3_) for all methods across different settings.

**Table 1:**
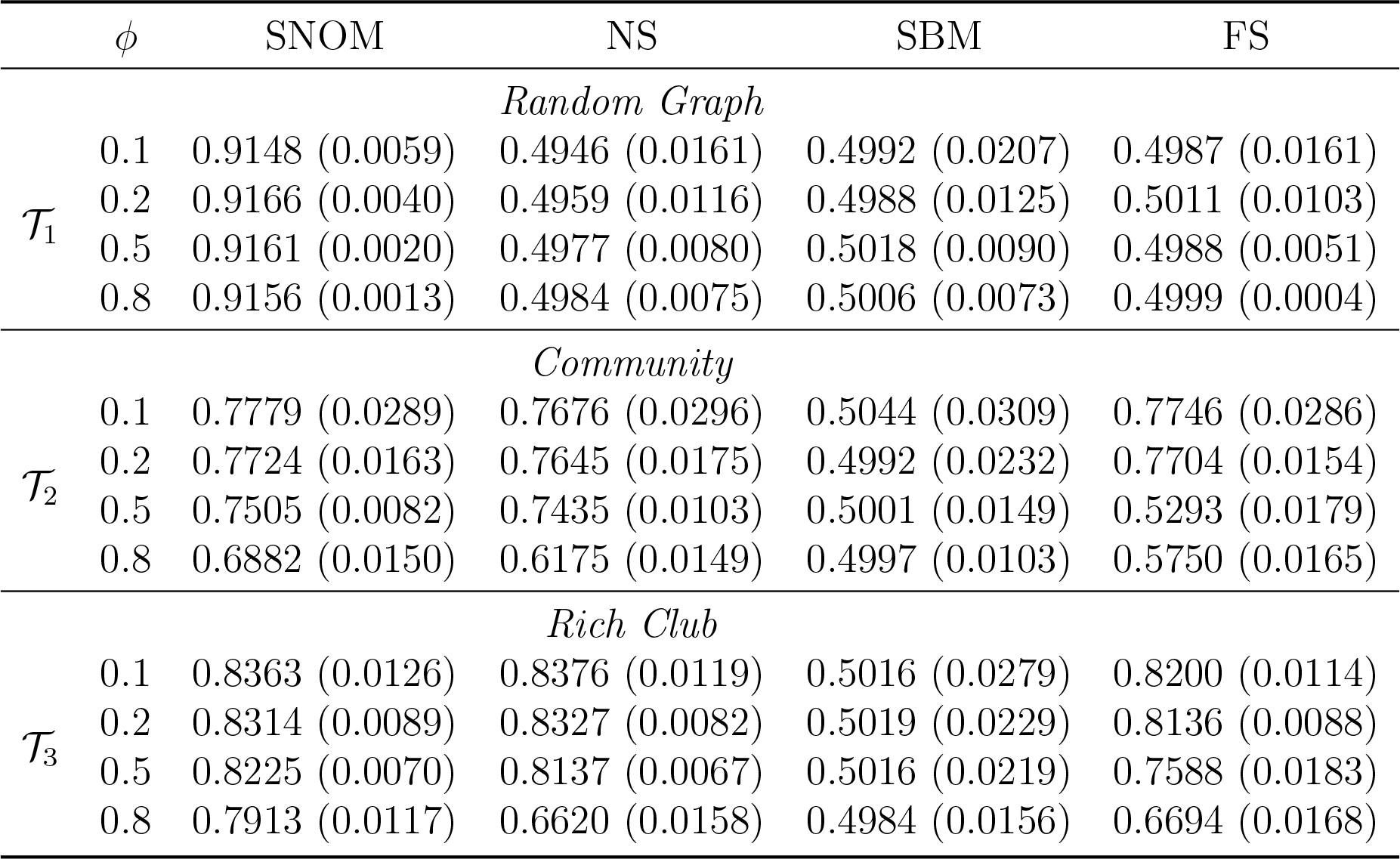
The results of latent contact prediction for all methods: the means and standard deviations across all simulated datasets of different settings.

### 3.3 Parameter estimation with validation data and pre-specified θ

In this section, we demonstrate the proposed method can provide an accurate estimation of ***β*** when *θ* can be pre-specified using the validation data. We let covariates *X*_1_ *∼* exp(1) and *X*_2_ *∼ N* (0, 1) and *β*_0_ = 0.5*, β*_1_ = 1*, β*_2_ = 1.5. The results in Table 2 show that both point and interval estimates are accurate and reliable when *θ* is correctly pre-specified. The results also indicate that our method with novel misclassified outcome specification (2) and expected likelihood (3) can provide an accurate estimation of the effects of exposure when validation data is provided. Therefore, the model can be readily applied when validation data is available and the parameter estimation procedure requires step one alone.

**Table 2:** The estimation results of regression coefficients with correctly pre-specified *θ*

|  |  | Para. | Est. | MSE | Coverage of 95% C.I. |
| --- | --- | --- | --- | --- | --- |
| $\theta = 0.05$ | n=50 | $\beta_0$ | 0.4959 | 0.0121 | 0.9660 |
| | | $\beta_1$ | 1.0073 | 0.0124 | 0.9650 |
| | | $\beta_2$ | 1.5064 | 0.0093 | 0.9730 |
| | n=150 | $\beta_0$ | 0.4997 | 0.0014 | 0.9480 |
| | | $\beta_1$ | 1.0000 | 0.0014 | 0.9640 |
| | | $\beta_2$ | 1.4993 | 0.0010 | 0.9620 |
| $\theta = 0.1$ | n=50 | $\beta_0$ | 0.5017 | 0.0107 | 0.9730 |
| | | $\beta_1$ | 0.9985 | 0.0111 | 0.9700 |
| | | $\beta_2$ | 1.4916 | 0.0078 | 0.9820 |
| | n=150 | $\beta_0$ | 0.5020 | 0.0012 | 0.9580 |
| | | $\beta_1$ | 0.9941 | 0.0013 | 0.9660 |
| | | $\beta_2$ | 1.4863 | 0.0010 | 0.9610 |
| $\theta = 0.15$ | n=50 | $\beta_0$ | 0.5077 | 0.0093 | 0.9830 |
| | | $\beta_1$ | 0.9874 | 0.0098 | 0.9700 |
| | | $\beta_2$ | 1.4660 | 0.0074 | 0.9680 |
| | n=150 | $\beta_0$ | 0.5095 | 0.0012 | 0.9700 |
| | | $\beta_1$ | 0.9809 | 0.0015 | 0.9490 |
| | | $\beta_2$ | 1.4599 | 0.0023 | 0.8510 |

## 4. Data Example

### 4.1 Example 1: Partially Observed Social Network

We apply the proposed method to a partially observed social network from a student cohort study for influenza research at the University of Maryland. One goal of this study is to investigate how environmental factors and biomarkers impact disease transmission between subjects with close contacts. The first phase of this study (year 2017) includes 75 undergraduate students age between 18 to 19 and 55 females and 18 males. The social network survey was conducted at the baseline, each study subject was asked to provide their four close contacts and 189 edges were recorded (reported case-contact connections). Clearly, many potential links are unreported which may misguide the results and yield biased estimates. Therefore, we are motivated to predict latent links and provide guidance to recover the latent links to better understand the disease transmission process.

We perform the link prediction using various methods and compare the performance based on the hold-out links (we artificially hide a proportion of edges beside the latent links from the observed data). The graph statistics used our method are determined by a variable selection procedure. For each link-hiding proportion, we repeat the analysis and prediction for 100 times and demonstrate the performance of each predictive model using the AUCs of averaged ROC curves in Table 3 and Figure 2.

**Figure 2:**
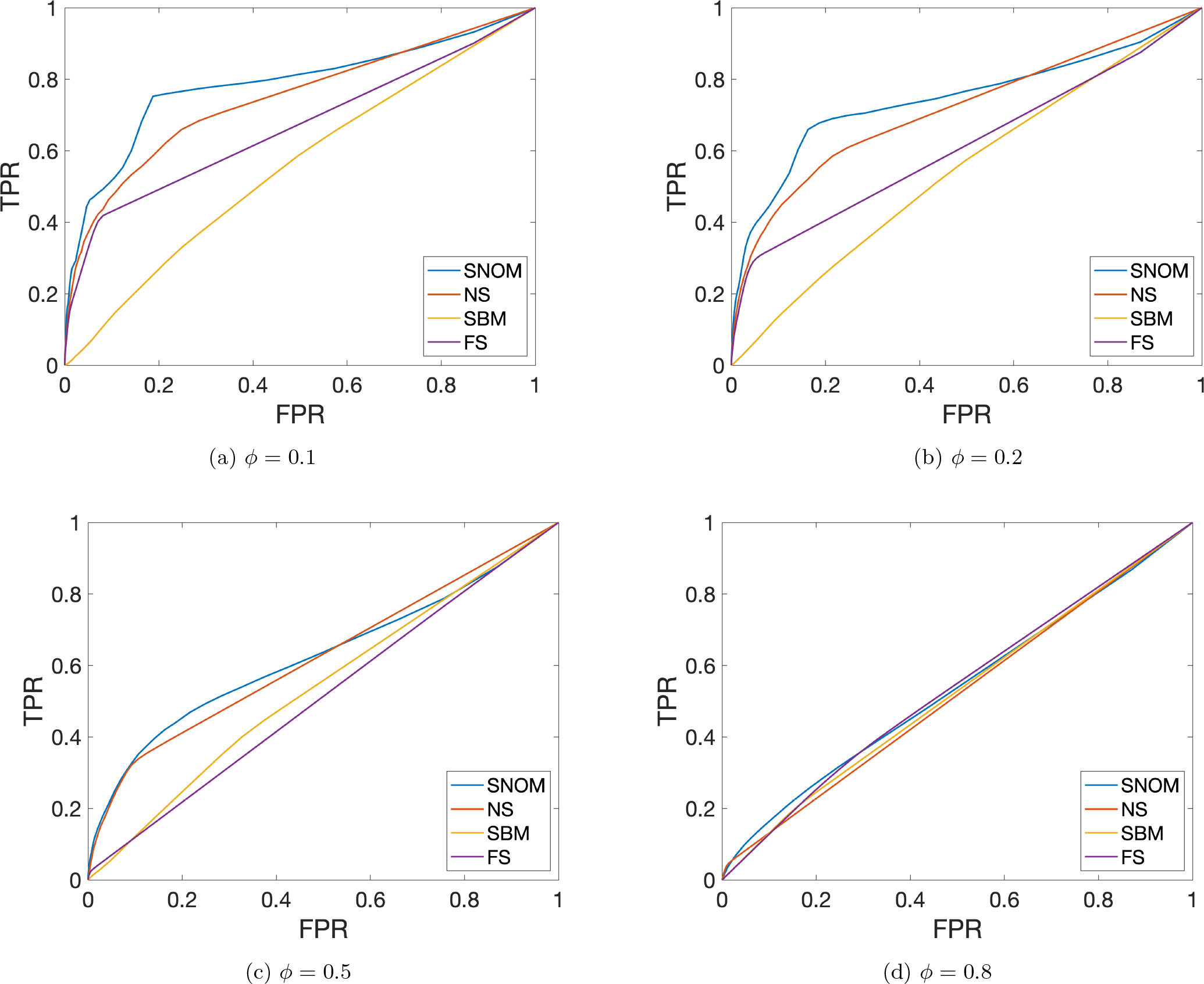
Averaged ROC curves for methods with different rates of hidden links in partially observed social networks

**Table 3:** Comparing AUCs of ROCs between methods for hold-out links in partially observed social networks

| $\phi$ | SNOM | NS | SBM | FS |
| --- | --- | --- | --- | --- |
| 0.1 | 0.7842 (0.0665) | 0.7460 (0.0617) | 0.5559 (0.0596) | 0.6660 (0.0548) |
| 0.2 | 0.7437 (0.0559) | 0.7150 (0.0387) | 0.5461 (0.0462) | 0.6124 (0.0412) |
| 0.5 | 0.6262 (0.0333) | 0.6227 (0.0280) | 0.5383 (0.0235) | 0.5128 (0.0151) |
| 0.8 | 0.5336 (0.0361) | 0.5172 (0.0126) | 0.5239 (0.0142) | 0.5357 (0.0226) |

Our method outperforms the competing methods when link hiding proportions are 0.1 and 0.2. Both the sensitivity and specificity of the predicted outcomes based on our methods are superior to the competing methods. In practice, the proportion of all non-connected links are latent links (i.e. the false negative rate) is around 0.1 and 0.2. Therefore, the proposed approach can be useful for many partially observed network data in epidemiological studies. When the unreported link rate increases to 0.5, the performance of our method is comparable to NS and superior to the other two methods. Last, all models fail when the rate of unreported links is 0.8 because the remaining information would not support proper predictive modeling. The results of this data example are well aligned with the results in the simulation section. The difference between our methods and competing methods of the data example is smaller than the social network with random graph structure (𝒯_1_) but greater than the social network with a very organized topological structure (𝒯_2_ and (𝒯_3_)). This may be explained by the fact that the real-world social network is composed of both organized topological patterns and randomness.

In addition, we apply our link prediction model to all observed links in the data example. In Figure 3, interestingly we note that our method predicts many latent links to be positive in the communities although the community structure is unknown prior to model estimation. It can be explained by that the ERGM can capture the network topological structures. The prediction seems to be reasonable because a group of students in college tends to take courses, eat, and relax in shared space although the validation data is not available and the ground truth can not be assessed. The predicted links will be used for further epidemiological investigation, for example, whether the dorm environmental factors can impact the transmission of infectious disease between contacted subjects.

**Figure 3:**
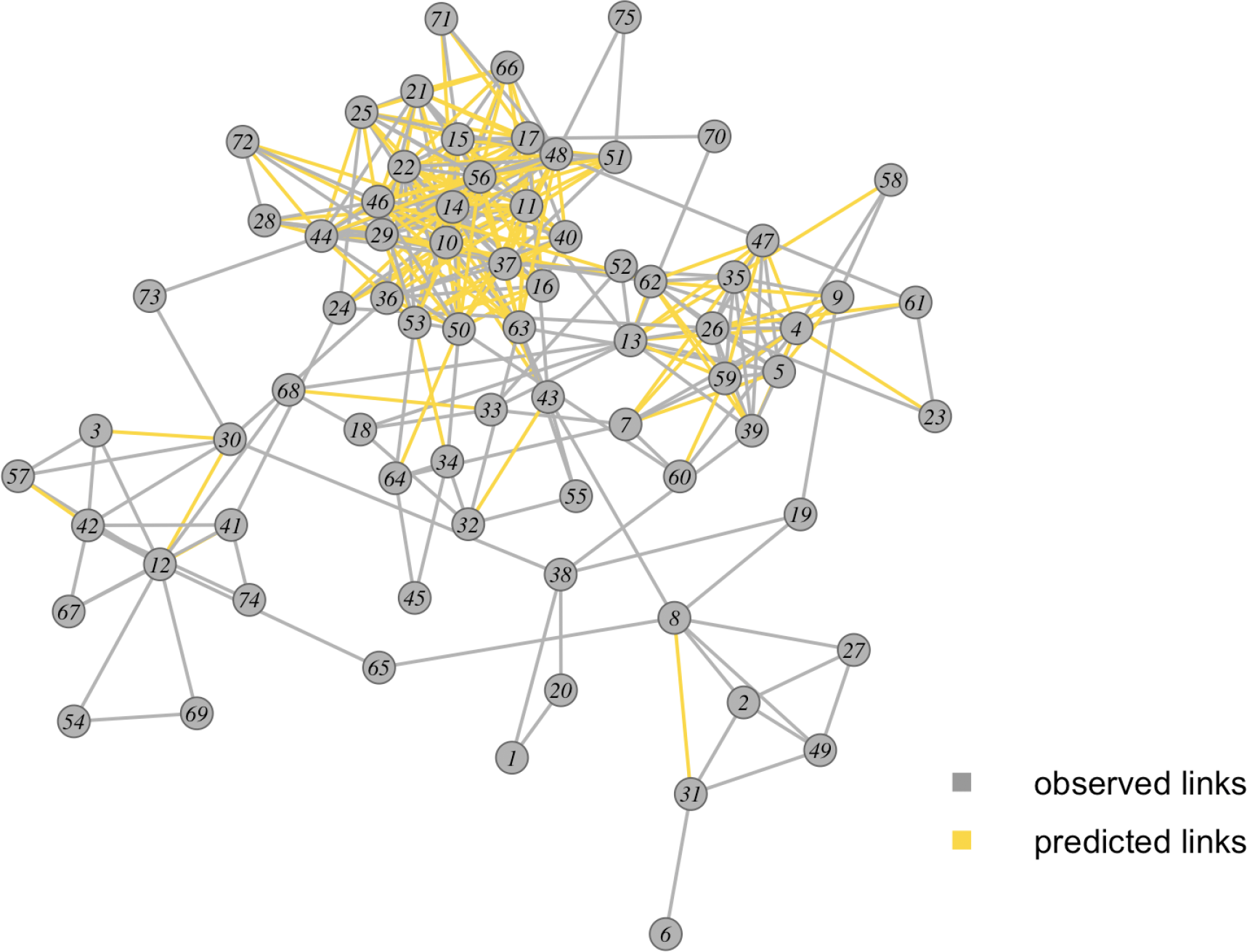
Predicted Social Network

### 4.2 Example 2: Incomplete Brain Connectivity Network

It has been documented that the graph properties and structures of brain networks largely resemble social networks (De Vico Fallani et al., 2014). In a brain network, each node represents a brain area and an edge/link represents the interactive relationship between a pair of brain areas (Rubinov and Sporns, 2010). The brain network is often thresholded to turn negative and weakly-connected edges into unconnected links (Van Wijk et al., 2010,Simpson et al., 2019). Although thresholding can remove false positive edges, it inevitably introduces false negative findings by setting links under detection limit as unconnected edges. Therefore, the thresholded brain network is considered a incomplete network. Our goal is recover the ‘true’ network based on the incomplete brain network data by identifying the false negative edges using our method. The brain network is built based on resting state fMRI data from 44 healthy subjects (age 36 *±* 11, 17 females and 27 males) from a previous study (Millman et al., 2019). The brain network includes 246 nodes defined by brain regions (Fan et al., 2016). The strengths of edges are measured by the temporal coherence between averaged time series from all brain regions/nodes (Chen et al., 2018, Chen et al., 2020). The group level (average) brain network characterizes the brain circuitry of healthy subjects at rest. Since the brain network is thresholded and many edges can be latent links due to detection limit, we apply the proposed method to the incomplete brain network and perform latent link prediction. We evaluate the performance of varied link prediction methods based on the artificially hold-out links. We generate 20 sets of hold-out links for each link-hiding proportion, and predict latent links based on the rest of edges in the observed brain network. The results of each predictive model using the AUCs of averaged ROC curves are displayed in Table 4 and Figure 4.

**Figure 4:**
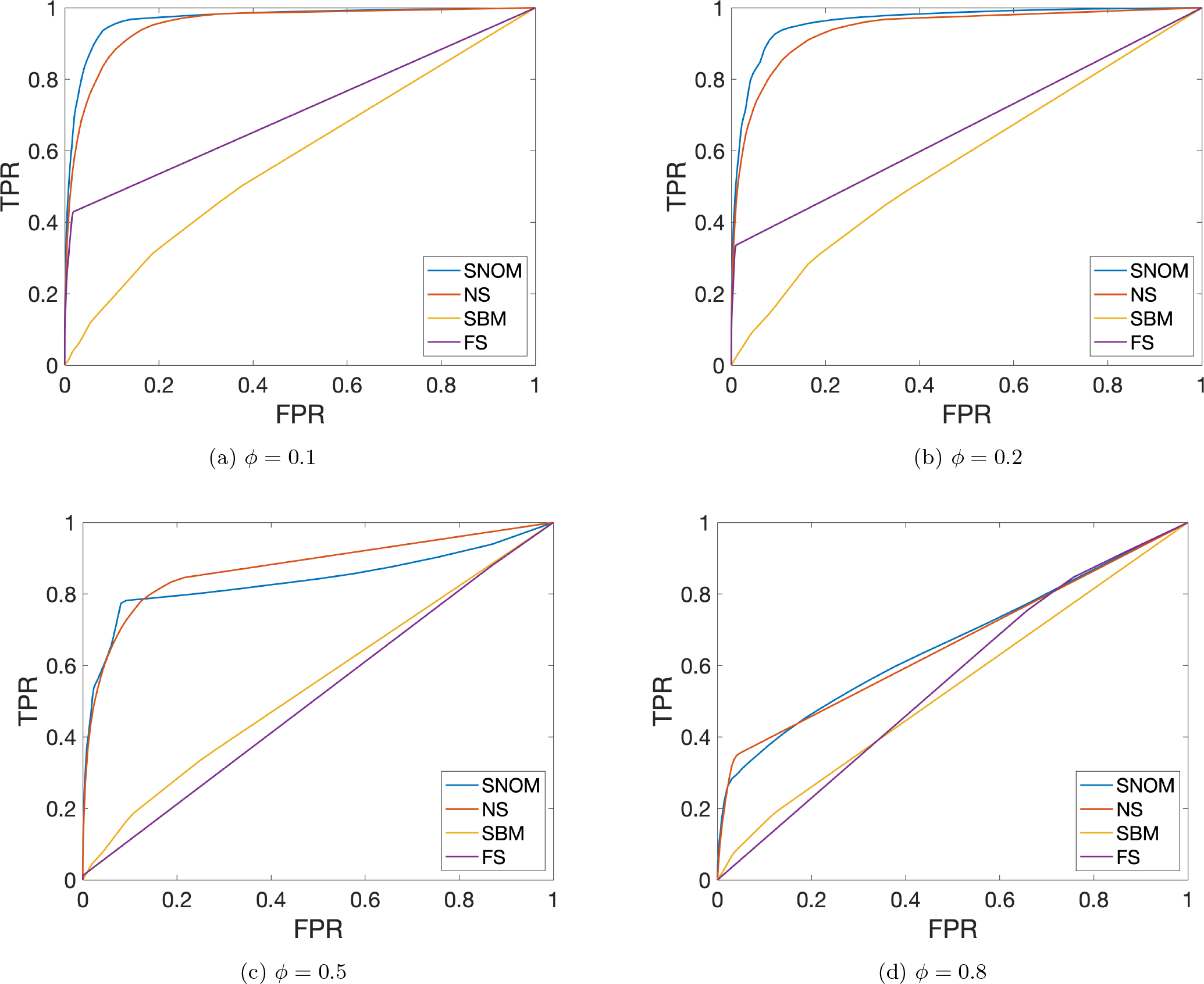
Averaged ROC curves for methods with different rates of hidden links in incomplete brain network

**Table 4:** Comparing AUCs of ROCs between methods for hold-out links in incomplete brain network

| $\phi$ | SNOM | NS | SBM | FS |
| --- | --- | --- | --- | --- |
| 0.1 | 0.9689 (0.0076) | 0.9529 (0.0104) | 0.5777 (0.0205) | 0.7073 (0.0196) |
| 0.2 | 0.9631 (0.0039) | 0.9405 (0.0079) | 0.5724 (0.0193) | 0.6638 (0.0122) |
| 0.5 | 0.8355 (0.0101) | 0.8719 (0.0098) | 0.5464 (0.0117) | 0.5126 (0.0119) |
| 0.8 | 0.6605 (0.0094) | 0.6568 (0.0150) | 0.5345 (0.0072) | 0.5523 (0.0134) |

Similarly to the social network example, our method SNOM outperforms the competing methods under low hiding rate (0.1 and 0.2). NS shows better performance than SBM and FS methods. For a higher hiding proportion 0.5,our method and NS method have similar performance and outperform the other two methods. When the hold-out rate increases to 0.8, no method can provide accurate prediction though NS method and our method outperform the other two methods. We note that the link prediction performance based on incomplete brain network (average AUC=97%) is generally better than the incomplete social network (average AUC=78%). This may be explained by the fact that social network data is collected based on subjective survey/questionnaire data which may include more noise than objectively observed neurophysiological signals of brain connectome network data.

## 5. Discussion

We develop a link prediction method for partially network data in the context of outcome misclassification and ERGM. Our emphasis is on “outcome/link prediction” which distinguishes our work from the prior work of binary outcome misclassification in aiming at “estimating effects of exposures” (Lyles et al., 2011). The predicted latent links provide a more accurate recovery of the full network data using only observed incomplete network data.

Our method complements the parametric existing social network model by allowing misclassified outcome data and unreported/latent links. We extend the basis of existing social network models from the likelihood to expected likelihood to account for the misclassified outcome variables. This framework is flexible to incorporate node and edge characteristics with the parametric model specification.

We also make several methodological contributions for the models of outcome misclassification for network analysis. The previous research has shown that validation data set is critical for parameter estimation for binary outcome misclassification adjustment. When validation data is available, the proposed method provides a new statistical model to account for the new (single-directional) misclassification mechanism (latent links) which yields both accurate outcome prediction and parameter estimation. However, the validation data for a partially observed network data is rarely available. Our theoretical proof and numerical results demonstrate that the lack of validation data has little impact on latent link prediction in partially observed social network data by leveraging the new optimization procedure. This conclusion of outcome prediction without validation data may be extended to the general binary outcome regression models beyond statistical network models with further investigations and experiments. The novel outcome misclassification specification for the single-directional misclassification is well suited for the efficient computation of expected likelihood.

We evaluate the proposed method using three network topological structures in the simulation study and two data examples from the undergraduate student cohort and a brain connectome study. Our method demonstrates improved and/or comparable performance with the existing machine learning and nonparametric methods in all scenarios. In practice, the complex network data including both social network and brain connectome network, presents combined patterns of organized topological structures and randomness. Our model provide accurate prediction of the latent links. In addition, our method is flexible and can incorporate information link-related multivariate covariates and latent network topological structures into network models. In summary, the proposed link prediction model can integrate information from the observed network and other covariates and effectively recover the full network based on incomplete network data.

## 6. Software

Software in the form of Matlab code, together with a sample input data set and complete documentation is available in https://github.com/qwu1221/LinkPred.

## Appendix

### A1. Numerical evaluation of the impact of θ estimation on link prediction by Thereom 1

To further validate theory 1, we perform numerical experiments to examine the impact of biased θ estimation on link prediction. We generate a social network with 50 nodes and sample edges (*y_ij_, i, j* = 1, …, 50) from a Bernoulli(π_ij_) distribution, where π_ij_ is linear combination of *X*_1_ ∼ exp(1) and *X*_2_ ∼ N (0, 1). We let observed edges w_ij_ are sampled from edges y_ij_ = 1 using another Bernoulli distribution with probability *ϕ_ij_* (*θ*). *θ*_0_ = 0.1 was used to generate the data.

We evaluate the impact of biased 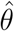 on c statistic by letting 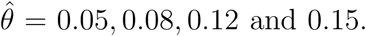 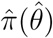 are calculated based on the step one optimization for all possible 1225 (50 × (50- 1)/2) links in the network. Next, the 749700 (1225× (1225-1)/2) pairs of links are used to count the concordant and discordant pairs. The AUC of ROC (which approximately equals to the c statistic) is determined by the proportion of concordant pairs. If for all 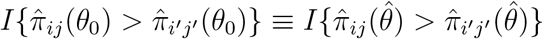, the proportion of concordant pairs is unchanged and the c statistic (AUC ROC) is not impacted by the estimation bias. Therefore, we can evaluate the impact of biased estimation of θ on the performance of link prediction by comparing the signs of 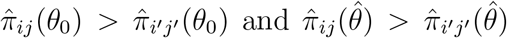. The simulation results in Table 1 indicate that less than 0.1% concordant and discordant pairs are affected by the biased θ^^^. Therefore, we conclude that moderately biased 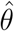 due to the lack of validation data set has little impact on the prediction of latent links for our method.

---

**Table A1:** The agreement of concordance and discordance metrics between

| | | $\theta_0 = 0.1$ | |
| --- | --- | --- | --- |
| | | $\hat{\pi}_{ij}(\theta_0) > \hat{\pi}_{i'j'}(\theta_0)$ | $\hat{\pi}_{ij}(\theta_0) < \hat{\pi}_{i'j'}(\theta_0)$ |
| $\hat{\theta} = 0.05$ | $\hat{\pi}_{ij}(\hat{\theta}) > \hat{\pi}_{i'j'}(\hat{\theta})$ | 384581 (0.5130) | 284 (3.7882e-04) |
| | $\hat{\pi}_{ij}(\hat{\theta}) < \hat{\pi}_{i'j'}(\hat{\theta})$ | 336 (4.4818e-04) | 364499 (0.4862) |
| $\hat{\theta} = 0.08$ | $\hat{\pi}_{ij}(\hat{\theta}) > \hat{\pi}_{i'j'}(\hat{\theta})$ | 384787 (0.5133) | 111 (1.7340e-04) |
| | $\hat{\pi}_{ij}(\hat{\theta}) < \hat{\pi}_{i'j'}(\hat{\theta})$ | 130 (1.4806e-04) | 364672 (0.4864) |
| $\hat{\theta} = 0.12$ | $\hat{\pi}_{ij}(\hat{\theta}) > \hat{\pi}_{i'j'}(\hat{\theta})$ | 384812 (0.5133) | 105 (1.4006e-04) |
| | $\hat{\pi}_{ij}(\hat{\theta}) < \hat{\pi}_{i'j'}(\hat{\theta})$ | 105 (1.4006e-04) | 364678 (0.4864) |
| $\hat{\theta} = 0.15$ | $\hat{\pi}_{ij}(\hat{\theta}) > \hat{\pi}_{i'j'}(\hat{\theta})$ | 384648 (0.5131) | 281 (3.7482e-04) |
| | $\hat{\pi}_{ij}(\hat{\theta}) < \hat{\pi}_{i'j'}(\hat{\theta})$ | 269 (3.5881e-04) | 364502 (0.4862) |

## References

Airoldi, E. M., Costa, T. B., and Chan, S. H. (2013). Stochastic blockmodel approximation of a graphon: Theory and consistent estimation. In Advances in Neural Information Processing Systems, pages 692–700.

Al Hasan, M., Chaoji, V., Salem, S., and Zaki, M. (2006). Link prediction using supervised learning. In SDM06: workshop on link analysis, counter-terrorism and security.

Austin, P. C. and Steyerberg, E. W. (2014). Graphical assessment of internal and external calibration of logistic regression models by using loess smoothers. Statistics in medicine 33, 517–535.

Carroll, R. J., Ruppert, D., Stefanski, L. A., and Crainiceanu, C. M. (2006). Measurement error in nonlinear models: a modern perspective. Chapman and Hall/CRC.

Chen, S., Bowman, F. D., and Xing, Y. (2020). Detecting and testing altered brain connectivity networks with k-partite network topology. Computational Statistics & Data Analysis 141, 109–122.

Chen, S., Xing, Y., Kang, J., Kochunov, P., and Hong, L. E. (2018). Bayesian modeling of dependence in brain connectivity data. Biostatistics.

Clauset, A., Moore, C., and Newman, M. E. (2008). Hierarchical structure and the prediction of missing links in networks. Nature 453, 98.

Copas, J. B. (1988). Binary regression models for contaminated data. Journal of the Royal Statistical Society: Series B (Methodological) 50, 225–253.

De Vico Fallani, F., Richiardi, J., Chavez, M., and Achard, S. (2014). Graph analysis of functional brain networks: practical issues in translational neuroscience. Philosophical Transactions of the Royal Society B: Biological Sciences 369, 20130521.

Fan, L., Li, H., Zhuo, J., Zhang, Y., Wang, J., Chen, L., Yang, Z., Chu, C., Xie, S., Laird, A. R., et al. (2016). The human brainnetome atlas: a new brain atlas based on connectional architecture. Cerebral cortex 26, 3508–3526.

Frank, O. and Strauss, D. (1986). Markov graphs. Journal of the american Statistical association 81, 832–842.

Goldenberg, A., Zheng, A. X., Fienberg, S. E., Airoldi, E. M., et al. (2010). A survey of statistical network models. Foundations and TrendsQR in Machine Learning 2, 129–233.

Harrell Jr, F. E. (2015). Regression modeling strategies: with applications to linear models, logistic and ordinal regression, and survival analysis. Springer.

Hoff, P. D., Raftery, A. E., and Handcock, M. S. (2002). Latent space approaches to social network analysis. Journal of the american Statistical association 97, 1090–1098.

Holland, P. W. and Leinhardt, S. (1981). An exponential family of probability distributions for directed graphs. Journal of the american Statistical association 76, 33–50.

Hunter, D. R., Handcock, M. S., Butts, C. T., Goodreau, S. M., and Morris, M. (2008). ergm: A package to fit, simulate and diagnose exponential-family models for networks. Journal of statistical software 24, nihpa54860.

Kingman, J. F. et al. (1978). Uses of exchangeability. The Annals of Probability 6, 183–197.

Li, K.-C., Duan, N., et al. (1989). Regression analysis under link violation. The Annals of Statistics 17, 1009–1052.

Lyles, R. H. and Lin, J. (2010). Sensitivity analysis for misclassification in logistic regression via likelihood methods and predictive value weighting. Statistics in medicine 29, 2297– 2309.

Lyles, R. H., Tang, L., Superak, H. M., King, C. C., Celentano, D. D., Lo, Y., and Sobel, J. D. (2011). Validation data-based adjustments for outcome misclassification in logistic regression: an illustration. Epidemiology (Cambridge, Mass.) 22, 589.

Magder, L. S. and Hughes, J. P. (1997). Logistic regression when the outcome is measured with uncertainty. American journal of epidemiology 146, 195–203.

Martínez, V., Berzal, F., and Cubero, J.-C. (2017). A survey of link prediction in complex networks. ACM Computing Surveys (CSUR) 49, 69.

Miller, K., Jordan, M. I., and Griffiths, T. L. (2009). Nonparametric latent feature models for link prediction. In Advances in neural information processing systems, pages 1276–1284.

Millman, Z. B., Gallagher, K., Demro, C., Schiffman, J., Reeves, G. M., Gold, J. M., Rouhakhtar, P. J. R., Fitzgerald, J., Andorko, N. D., Redman, S., et al. (2019). Evidence of reward system dysfunction in youth at clinical high-risk for psychosis from two event-related fmri paradigms. Schizophrenia research.

Neuhaus, J. M. (1999). Bias and efficiency loss due to misclassified responses in binary regression. Biometrika 86, 843–855.

O’Madadhain, J., Hutchins, J., and Smyth, P. (2005). Prediction and ranking algorithms for event-based network data. ACM SIGKDD explorations newsletter 7, 23–30.

Reilly, M. and Pepe, M. S. (1995). A mean score method for missing and auxiliary covariate data in regression models. Biometrika 82, 299–314.

Rhodes, C. and Jones, P. (2009). Inferring missing links in partially observed social networks. Journal of the operational research society 60, 1373–1383.

Robins, G., Pattison, P., Kalish, Y., and Lusher, D. (2007). An introduction to exponential random graph (p*) models for social networks. Social networks 29, 173–191.

Rubinov, M. and Sporns, O. (2010). Complex network measures of brain connectivity: uses and interpretations. Neuroimage 52, 1059–1069.

Shojaie, A. (2013). Link prediction in biological networks using multi-mode exponential random graph models. In 11th Workshop on Mining and Learning with Graphs, pages 987–991.

Simpson, S. L., Bahrami, M., and Laurienti, P. J. (2019). A mixed-modeling framework for analyzing multitask whole-brain network data. Network Neuroscience 3, 307–324.

Simpson, S. L., Hayasaka, S., and Laurienti, P. J. (2011). Exponential random graph modeling for complex brain networks. PloS one 6, e20039.

Simpson, S. L. and Laurienti, P. J. (2015). A two-part mixed-effects modeling framework for analyzing whole-brain network data. NeuroImage 113, 310–319.

Snijders, T. A. and Van Duijn, M. A. (2002). Conditional maximum likelihood estimation under various specifications of exponential random graph models. Contributions to social network analysis, information theory, and other topics in statistics pages 117–134.

Stattner, E. and Vidot, N. (2011). Social network analysis in epidemiology: Current trends and perspectives. In 2011 Fifth International Conference on Research Challenges in Information Science, pages 1–11. IEEE.

Strauss, D. and Ikeda, M. (1990). Pseudolikelihood estimation for social networks. Journal of the American statistical association 85, 204–212.

Van Wijk, B. C., Stam, C. J., and Daffertshofer, A. (2010). Comparing brain networks of different size and connectivity density using graph theory. PloS one 5, e13701.

Wasserman, S. and Pattison, P. (1996). Logit models and logistic regressions for social networks: I. an introduction to markov graphs andp. Psychometrika 61, 401–425.

Zhang, M. and Chen, Y. (2018). Link prediction based on graph neural networks. In Advances in Neural Information Processing Systems, pages 5165–5175.

Zhang, Y., Levina, E., and Zhu, J. (2017). Estimating network edge probabilities by neighbourhood smoothing. Biometrika 104, 771–783.

Zhao, Y., Wu, Y.-J., Levina, E., and Zhu, J. (2017). Link prediction for partially observed networks. Journal of Computational and Graphical Statistics 26, 725–733.

Zhou, S. and Mondragón, R. J. (2004). The rich-club phenomenon in the internet topology. IEEE Communications Letters 8, 180–182.

